# Oligodendrocyte subtype diversity underlines clinical progression in Parkinson’s disease

**DOI:** 10.64898/2026.05.15.724963

**Authors:** Daniela Mirzac, Nils Schröter, Yann Decker, Martin B. Glaser, Matthias Hülser, Svenja L. Kreis, Heiko J. Luhmann, Viviane Almeida, Jenny Blech, Sebastian Kunz, Matthias Klein, Tobias Bopp, Michael T. Heneka, Philip L. De Jager, Joachim Oertel, Gabriel Gonzalez-Escamilla, Sergiu Groppa

**Affiliations:** Department of Neurology, Saarland University Hospital and Saarland University; Homburg, Germany; Neuroimaging Center, University Medical Center of the Johannes Gutenberg University Mainz; Mainz, Germany; Department of Neurosurgery, University Medical Center, Johannes Gutenberg University Mainz; Mainz, Germany; Institute of Physiology, University Medical Center, Johannes Gutenberg University Mainz; Mainz, Germany; Institute of Immunology, Center for Immunotherapy (FZI), University Medical Center, Johannes Gutenberg University Mainz; Mainz, Germany; Luxembourg Centre for Systems Biomedicine; Belvaux, Luxembourg; Center for Translational & Computational Neuroimmunology, Department of Neurology, Taub Institute for Research on Alzheimer’s Disease and the Aging Brain, Columbia University Irving Medical Center; New York, NY, USA

## Abstract

Despite growing evidence for glial involvement in Parkinson’s disease, oligodendrocyte dysfunction remains poorly defined. To address this gap, we compared single-cell RNA sequencing from a mouse model of α-synuclein aggregation pathology with fresh human brain tissue from deep brain stimulation surgery to build a cross-species framework of disease progression. In total, we profiled over 200,000 cortical transcriptomes, including 55,000 oligodendrocytes. Early disease in mice was characterized by inflammatory activation, while advanced stages in both species converged on metabolic dysfunction, including impaired ribosomal output, chaperone stress responses, ubiquitination deficits, and lysosomal perturbation. In patients, APLP1 was upregulated and correlated with clinical disease progression and increased levodopa demand, linking α-synuclein spread in oligodendrocytes to disease severity. APP and CNTN pathways emerged as key signalling axes, with CNTN reflecting weakened reparative communication and reduced resilience. Together, these findings define oligodendrocyte subtype dynamics as shared and clinically relevant features of PD progression.

## INTRODUCTION

Parkinson’s disease (PD), the second most common neurodegenerative disorder, is characterized by progressive loss of dopaminergic neurons alongside the accumulation of misfolded α-synuclein in Lewy bodies, not only within neurons but also in other cell types [1]. Clinically, PD presents with motor symptoms driven by dysfunction in basal ganglia–thalamo–cortical circuits that converge on the primary motor and premotor cortex [2]. Concurrently, functional and structural alterations in cortical regions contribute to cognitive decline, impaired sensorimotor integration, and the emergence of non-motor symptoms [3]. Consistent with this, the cortex exhibits widespread α-synuclein pathology and disrupted network activity [4], reflecting altered neuronal–glial communication.

In this setting, oligodendrocytes appear particularly affected. The intercellular transfer of misfolded α-synuclein from neurons to oligodendrocytes likely contributes to the prominent inclusions observed in these cells [5]. Recent single-cell and single-nucleus transcriptomic studies of the human midbrain, substantia nigra, and cortex suggest that oligodendrocytes are among the most affected cell types in PD [6–12]. Reported transcriptional alterations include heightened stress and inflammatory responses [9, 10, 12], activation of the unfolded protein response [6] and diminished myelination capacity [6, 9].

By producing mediators such as CD74, C1QC, C1QB, FOS, and JUN, oligodendrocytes emerge as key components amplifying chronic neuroinflammatory cascades leading to neurodegeneration [13]. An important component for the contribution of the oligodendrocytes to neuroinflammation in PD is their interactions with microglia, astrocytes, and infiltrating immune cells, which alters their signalling profiles [14]. Despite these observations, the precise molecular mechanisms through which oligodendrocytes contribute to PD pathogenesis remain insufficiently resolved.

To address this gap and delineate the role of oligodendrocytes in PD, we analysed about 200,000 high-quality single-cell transcriptomes of fresh cortical samples from human patients and parkinsonian mouse model. The isolated ∼55,000 oligodendrocytes across species and disease stages resolved in transcriptionally distinct subtypes with individual functional signatures. We provide subtype-specific biological relevance at an early and late-disease stage in a mouse model and integrate the main findings within the human context. Using clinical correlation analyses, we linked these transcriptomic shifts to disease severity and levodopa treatment demand, both reflecting motor progression and dopaminergic therapeutic burden as clinically grounded proxies of disease trajectory. These proxies correlated with APLP1 upregulation in patients, linking α-synuclein spread in oligodendrocytes to disease severity. In parallel, APP and CNTN pathways emerged as key signalling axes, with CNTN depletion reflecting weakened reparative communication and reduced resilience. Together, these findings define oligodendrocyte subtype diversity in PD and implicate their transcriptional reprogramming in disease progression.

## RESULTS

### Oligodendrocyte dynamics across early and advanced stages in the mouse model of PD

To investigate oligodendrocyte subtype diversity across disease progression in PD, we employed a well-established mouse model based on α-synuclein aggregation–associated pathology [15]. Brain samples were collected from mice at defined early and advanced stages of the disease. For the early-stage, mice were analysed at four weeks after human A53T α-synuclein overexpression in the substantia nigra (Fig. 1 A, see Methods), a time point at which no overt motor impairment was observed despite an approximate 45% loss of dopaminergic neurons [15]. In contrast, for the advanced stage, mice were sacrificed eight weeks post-injection, when dopaminergic neuron loss reached ∼82% and was accompanied by detectable motor deficits [15].

**Figure 1.**
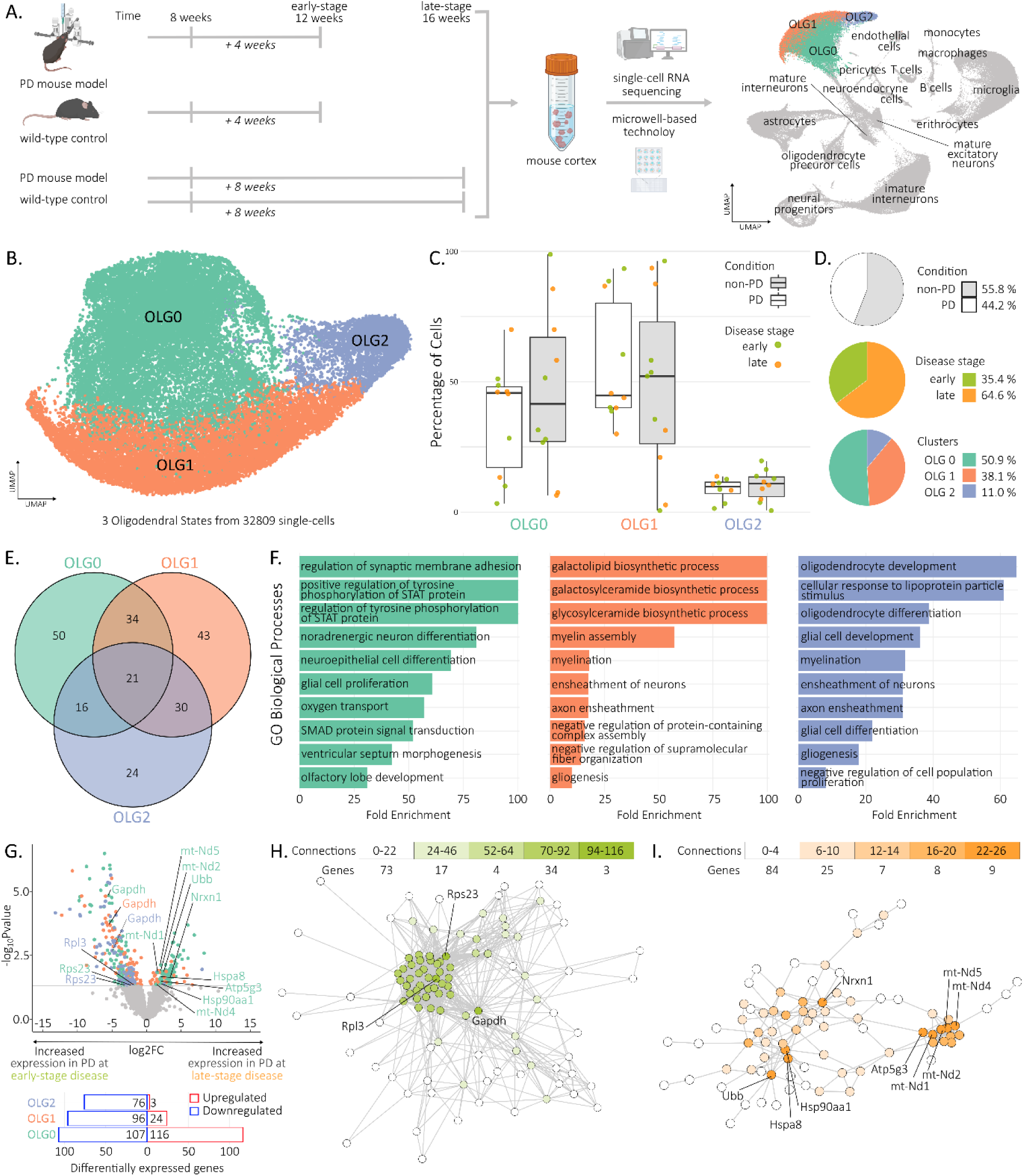
Subcellular diversity of oligodendrocytes in mice. (A) Experimental design on mice. (B) UMAP visualization of mouse oligodendrocyte subtypes, identifying three clusters based on marker gene expression and pathway enrichment. (C) Box-and-whisker plots of subtype proportions per mouse; grey: non-PD control mice; white: PD parkinsonian mice; green: four weeks; orange: eight weeks. (D) Pie charts of the distribution of cells by condition (non-PD and PD), disease stage (four weeks and eight weeks), subtypes (three clusters). (E) Venn diagram of shared and subtype-specific marker genes. (F) Top ten enriched ‘Gene Ontology Biological Process’ pathways for each subtype, ranked by enrichment fold and coloured by oligodendrocyte subtype. (G) Volcano plots of differentially expressed genes (DEG) comparing PD and non-PD mice within oligodendrocyte subtypes; significant genes are coloured by subtype and hub genes are labelled. Bar plots indicate numbers of up- and downregulated genes per subtype. (H,I) Protein–protein interaction network of upregulated DEGs at four weeks (H) and eight weeks (I). Network coloured by connectivity degree. Bar plot shows node counts. Top hub genes with most connections are labelled. UMAP Uniform Manifold Approximation and Projection; PD parkinsonian mouse model; non-PD control mice; OLG0-2 oligodendrocyte subtype 0-2.

We generated single-cell transcriptomic profiles from mouse cortical samples for 138,812 viable cells (Fig. 1 A), out of which 32,809 mouse oligodendrocytes were grouped in three sub-cellular types (Fig. 1 B, see Methods). Cells were evenly distributed across conditions, with 44.2% derived from the parkinsonian mouse model and 55.8% from controls, reflecting the initial cohort sizes (20 vs 22 mice; Fig. 1 C,D). In terms of disease progression, 35.4% of cells corresponded to the early-stage and 64.6% to the late-stage (Fig. 1 C,D). Oligodendrocyte subtype 0 comprised 50.9% of cells, with the remainder distributed between subtype 1 (38.1%) and subtype 2 (11.0%) (Fig. 1 B,C). We evaluated the individual markers of each subtype (Fig. 1 E), and their underlying dysregulated biological pathways (Fig. 1 F, see Methods). Importantly, while each subtype exhibited both shared and unique markers, they converged on common biological processes related to “myelination”, “glial cell proliferation and development”, and “axon ensheathment” (Fig. 1 F). This highlights extensive overlap among mouse oligodendrocytes, indicating limited subtype specialization and a largely homogeneous cellular state.

For analyses of disease progression specific states, we analysed transcriptomic alterations between early and late disease stages (Fig. S1 A). Early-stage mice showed a higher number of upregulated genes (Fig. S1 B), indicating that oligodendrocyte transcriptional alterations emerge early in the disease course, preceding the symptoms onset. Dysregulated genes showed minimal overlap across time points and oligodendrocyte subtypes (Fig. S1 C), with no evidence for a coordinated shift from early-stage upregulation to late-stage downregulation, or vice versa, at the gene level at this particular stage. (Fig. S1 D).

To identifying stage-specific biological processes, we performed network analyses of enriched and depleted pathways. Early-stage disease was marked by immune and inflammatory activation alongside impaired mitochondrial and oxidative metabolism (Fig. S1 E-G, left panels). In contrast, late-stage disease showed relative enrichment of energy metabolism, synaptic organization, and neurotransmission, accompanied by depletion of pathways related to neuronal support, including neuronal differentiation, axonogenesis, and guidance (Fig. S1 E–G, right panels). Together, these patterns indicate a shift from early inflammatory involvement to late-stage loss of oligodendrocyte-mediated neuronal support.

When contrasting differentially expressed genes between time points, we applied a multivariable model correcting for age and batch effects. This approach isolated changes specifically associated with disease stage, and revealed a similar pattern of dysregulation emerging. There are more genes involved at the early-stage compared to the latter stage (Fig. 1 F). The hub of upregulated genes at the early-stage, including Rsp23, Rpl3 and Gapdh (Fig. 1 G), is related to ribosome structure and protein translation [16], and cellular metabolism [17]. Conversely, at the late-stage disease there are two hubs of upregulated genes: mitochondrial and heat-shock proteins (Fig. 1 H). Heat shock proteins function as both sensors and effectors of proteotoxic stress by normally restraining transcription factors, which become activated when misfolded proteins accumulate [18]. This activation induces heat shock protein expression to restore proteostasis, and may account for the heat shock response observed in oligodendrocytes and other glial cells in PD [12].

To further investigate the alterations occurring later in the disease course affected by the α-synuclein propagation and emerging neurodegeneration, and bridge the translational gap from mouse models to clinical progression and human disease pathology we subsequently focused our analyses on human tissue.

### Oligodendrocyte subtype diversity in PD

To uncover the human oligodendrocyte subtype diversity, we analysed dorsolateral prefrontal cortex (DLPFC) samples from individuals with PD. In total, we profiled single-cells from 11 individuals with PD (mean age: 58.82 ± 13.86 years; disease duration: 9.73 ± 4.88 years) and contrasted them against 5 non-PD patients (mean age: 57.8 ± 16.2 years; disease duration: 13.4 ± 5.6 years) (Fig. 2 A, left panel; Supplementary table 1). Across all samples, we obtained transcriptomic profiles for 34,125 genes and 59,224 high-quality cells after filtering (see Methods, Fig. S2 A) [19]. Per individual, an average of 3,701 cells and 2,125 genes were analysed and subsequently resolved in 12 major cell-types manually annotated using canonical markers (Fig. 2 A, middle panel) [19, 20]. We next isolated ∼23,000 cells assigned to the oligodendrocyte cluster, defined by expression of canonical markers including MBP [21], PLP1 [22], MAG [23], and MOG [24]. Transcriptional profiling resolved seven distinct oligodendrocyte subtypes (Fig. 2 B). Consistent with previous reports [25], human oligodendrocytes retained transcriptional plasticity at later maturation stages, co-expressing shared and little subtype-specific markers (Fig. S2 B).

**Figure 2.**
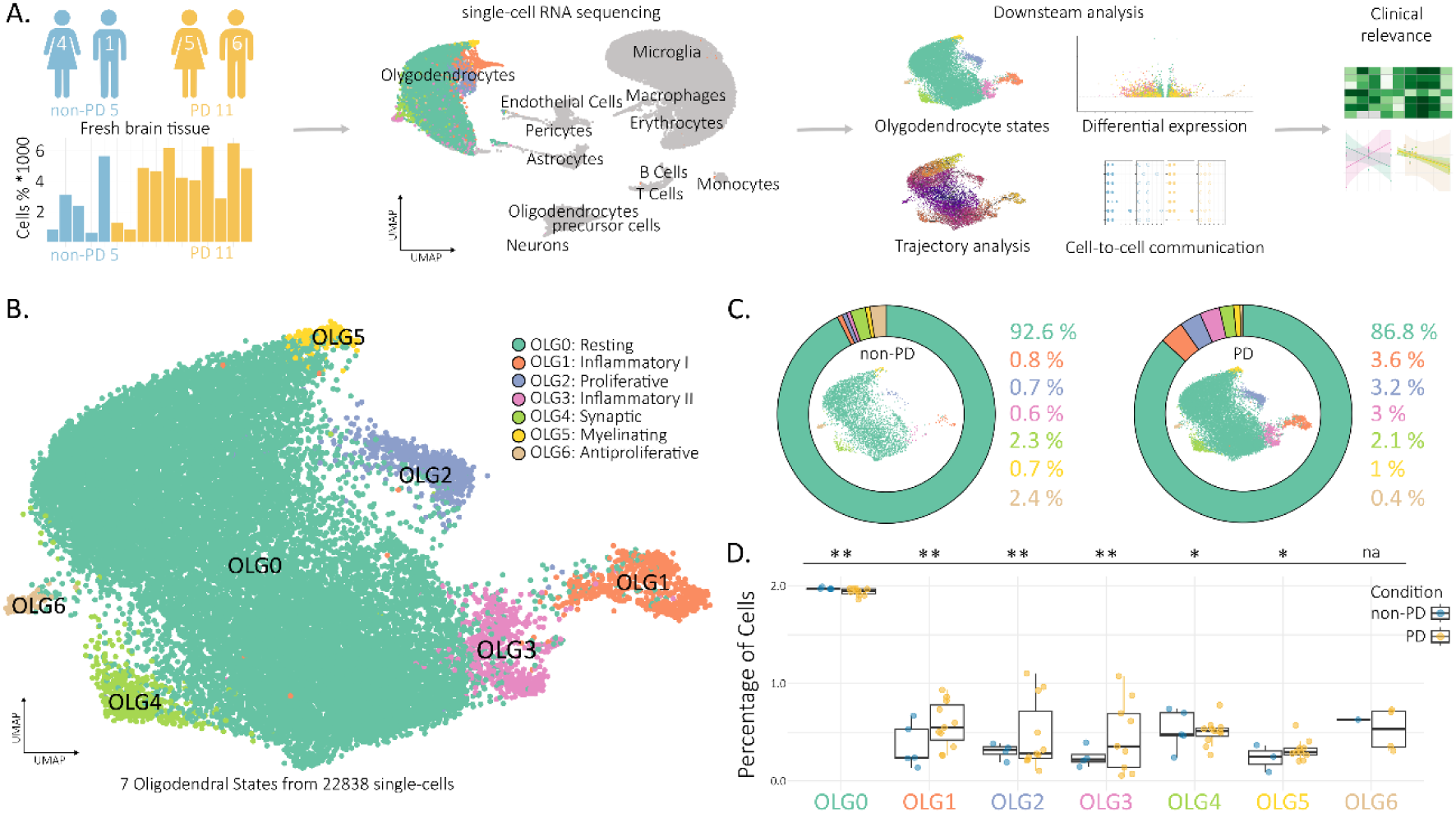
Subcellular diversity of oligodendrocytes in patients. (A) Experimental design on patients: 16 fresh human dorsolateral prefrontal cortex samples, including 11 from individuals with Parkinson’s disease (PD; yellow) and 5 from non-PD controls (blue). Bar plot shows the number of cells per individual included in the main analysis. Uniform Manifold Approximation and Projection (UMAP) embedding of 12 major cell types; oligodendrocytes are coloured by subcellular types. Summary of the downstream analytical workflow. (B) UMAP of oligodendrocyte subtypes. Seven clusters identified based on marker gene expression and pathway enrichment. (C) UMAP of oligodendrocyte subtypes split by cohort. Bar plots show subtype proportions per cohort, colored by subtype. Percentages are shown to the right of each plot. (D) Box-and-whisker plot of subtype proportions per individual. Blue: non-PD; yellow: PD. Statistical comparison performed using a linear mixed-effects model; P-values FDR-corrected. PD Parkinson’s disease; UMAP Uniform Manifold Approximation and Projection; OLG0-6 oligodendrocyte subtype 0-6.

In contrast to mice, human oligodendrocytes showed greater heterogeneity, resolving into multiple transcriptionally distinct subtypes. Subtypes 1 and 3 were defined by inflammatory markers (e.g., CD74, CCL3, C1QA; SPARCL1, CCL4) and enriched inflammatory pathways (Fig. S2, S3 A,C,E), while subtype 2 displayed FOS/JUN-driven signatures linked to proliferation, partially overlapping with inflammatory states (Fig. S2 B). Subtype 6 was associated with TGF-β1 regulation (Fig. S3 A,H). While subtype 5 (TUBB2B, TNFRSF21, S100A1) was enriched for myelination and axonal support (Fig. S2 B; S3 A,G), subtype 4 (CNTN1, FXYD6, KCNK10) displayed intracellular transport and synaptic functions (Fig. S2 B; Fig. S3 A,F). Subtype 0 showed low marker expression overall but enrichment in protein metabolism, consistent with a resting population (Fig. S2 C,D; S3 A,B). Although some marker overlap was observed between clusters, expression levels were generally low (Fig. S2 E), indicating that subtype identity is better defined by relative expression patterns and pathway context rather than distinct marker exclusivity.

The distribution of human oligodendrocyte subtypes across PD and non-PD cohorts revealed a higher overall cell density in PD, reflecting the larger number of cells and samples in this group (Fig. 2 C; S2 A). The predominance of the resting subtype 0 in non-PD individuals (92.6%) compared to PD (86.8%), hints at oligodendroglia disease related re-assignment. To this end, subtypes 1 to 3 (inflammatory and proliferative) were more prevalent in PD, with subtype 1 increasing from 0.8% to 3.6%, subtype 2 from 0.7% to 3.2%, and subtype 3 from 0.6% to 3.0% (Fig. 2 C). This corroborates a shift from the resting subtype 0 toward disease-associated transcriptional states. This shift was particularly evident in the density plot (Fig. S2 A), where subtype 1 to 3 cells were sparsely represented in non-PD, yet densely clustered in PD samples. Linear mixed-effects modelling confirmed these compositional changes (Fig. 2 D). Resting subtype 0 was significantly reduced in PD (-6.53%, q = 0.004), while inflammatory and proliferative subtypes 1 (+1.72%), 2 (+2.12%), and 3 (+2.24%) were significantly increased (q = 0.004). Smaller but significant differences were also observed for subtype 4 (-0.32%, q = 0.019) and subtype 5 (+0.38%, q = 0.019).

### Stage-dependent trajectories of oligodendrocyte subtypes and associated gene markers in PD

To further explore the transcriptional landscape and dynamic state transitions of human oligodendrocytes and their distinct gene signatures in PD, we performed pseudotime trajectory analysis using Monocle 3 (Fig. 3 A) [26]. The trajectory was anchored at the centre of subtype 0, where cell densities were comparable across PD and non-PD cohorts (Fig. S3 A), allowing for an unbiased starting point. The identified seven subtypes in pseudotime progression reveal a continuum of three major transcriptional stages (Fig. 3 C). First pseudotime stage encompasses cells from the proliferative subtype (OLG2), second stage includes inflammatory (OLG3) and myelinating (OLG5) subtypes, and third stage consists primarily of inflammatory (OLG1) and synaptic (OLG4) subtypes. The anti-proliferative subtype (OLG6) was not prominently displayed due to lower cell count compared to other subtypes. The resting subtype, due to its abundance and broad transcriptomic profile, had cells distributed throughout the continuum. The branching architecture of the pseudotime tree (Fig. 3 B) reveals a clear root-to-leaf structure, with each terminal branch dominated by a distinct oligodendrocyte subtype. This architecture supports the interpretation of subtypes as discrete, yet transcriptionally connected, states along a continuous trajectory.

**Figure 3.**
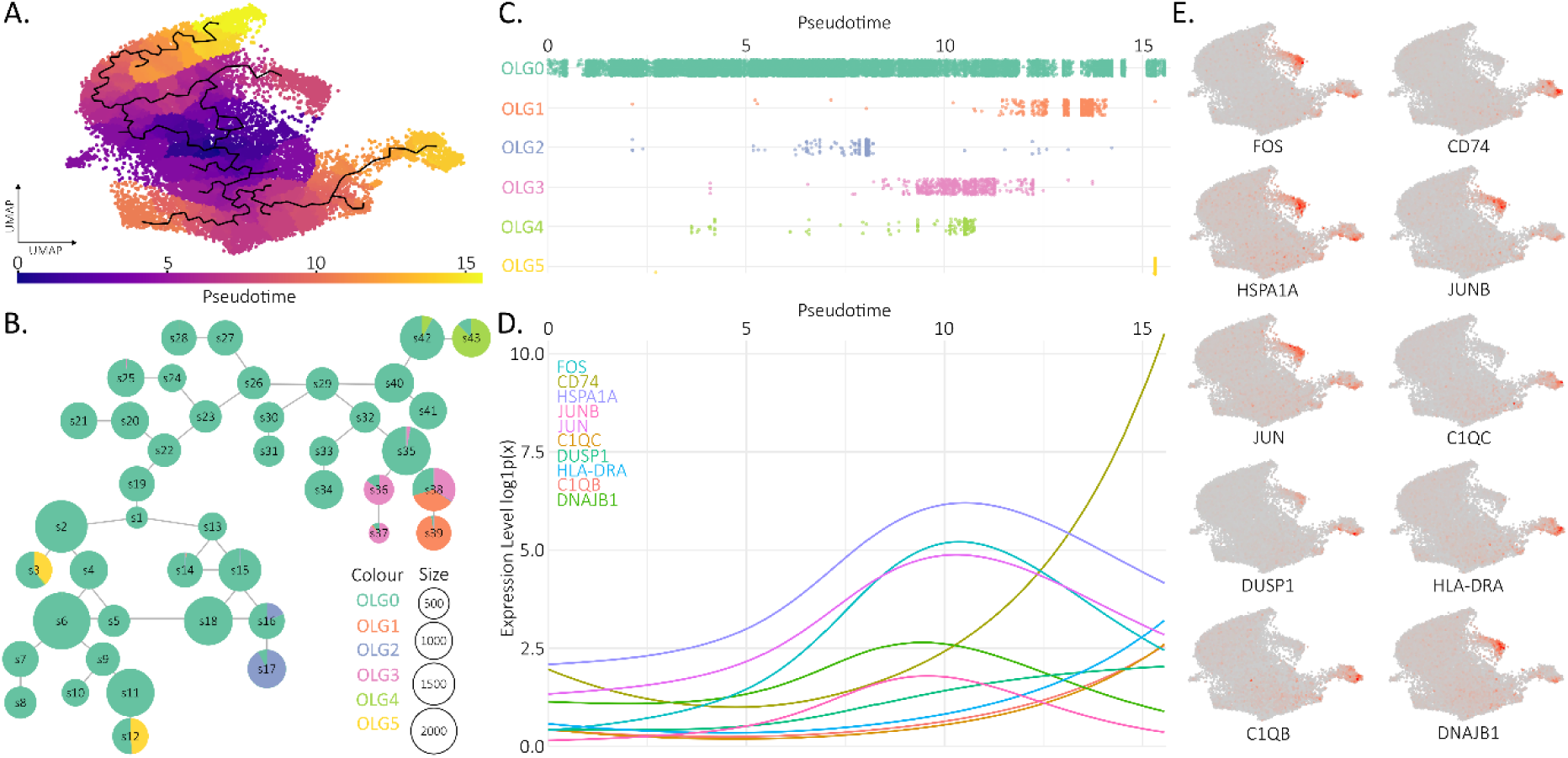
Inferred pseudotime trajectory of oligodendrocyte subtypes. (A) UMAP visualization of oligodendrocyte subtypes coloured by inferred pseudotime. (B) Trajectory tree plot showing identified nodes (root, branches, leaves); non-branching nodes were collapsed. Nodes are represented as pie charts, coloured by subtype composition. Pie size reflects the number of cells assigned to each node. (C) Dot plot showing distribution of cells across pseudotime, separated by oligodendrocyte subtype. (D) Pseudotime expression dynamics of the top 10 differentially expressed genes along the trajectory. Expression values were normalised as natural logarithm of (1 + x). (E) Feature plots of the top 10 differentially expressed genes across pseudotime. UMAP Uniform Manifold Approximation and Projection; OLG0-5 oligodendrocyte subtype 0-5; S Trajectory node reference.

We then examined the top ten genes differentially expressed along the pseudotime. First pseudotime stage, predominantly represented by the proliferative subtype, was defined mainly by five genes: FOS, JUNB, JUN, DNAJB1, and HSPA1A. These genes showed a sharp increase in expression coinciding with the abundance of respective cells (Fig. 3 D,E). Since the proliferative subtype is predominantly found in PD where a reactive, microglia-rich environment is present [12, 27], we interpret this transcriptional activation as a response to local inflammation. The early emergence of this state further suggests that oligodendrocytes actively contribute to the initiation of neuroinflammatory processes, as we elaborated in our previous work [12, 27]. In vitro studies further support this, showing that IL-1β released by activated microglia induces rapid upregulation of immediate early genes such as c-Fos and c-Jun in mixed glial cultures containing oligodendrocytes [28].

The second pseudotime stage includes inflammatory II and myelinating subtypes, did not feature prominently among the top genes due to overlapping expression patterns. In contrast, the third pseudotime stage, mainly represented by inflammatory I subtype, was defined by elevated expression of CD74, C1QC, C1QB, and HLA-DRA (Fig. 3 D,E).

Overall, these patterns highlight distinct gene expression shifts that mark different oligodendrocyte states along the trajectory, consistent transcriptional differences. Building on this, we next identified genes altered specifically in PD.

### Subtype-specific oligodendrocyte transcriptomic alterations in PD versus non-PD patients

To examine transcriptomic alterations in PD across oligodendrocyte subtypes, we performed pseudobulk differential expression analysis resulting in differentially expressed genes (DEGs; Fig. 4 A; Fig. S4 A,D.G,J,M,P,S). As expected, subtypes with more cells yielded a greater number of DEGs (Fig. 4B). In the resting subtype, genes were almost evenly split between up-and downregulation. In contrast, several subtypes showed a predominance of downregulated genes in PD. Only the anti-proliferative subtype displayed mostly upregulated genes and a single enriched pathway (i.e., ‘inflammatory response’ (Fig. 4 B,C; Fig. S4 S,T,U).

**Figure 4.**
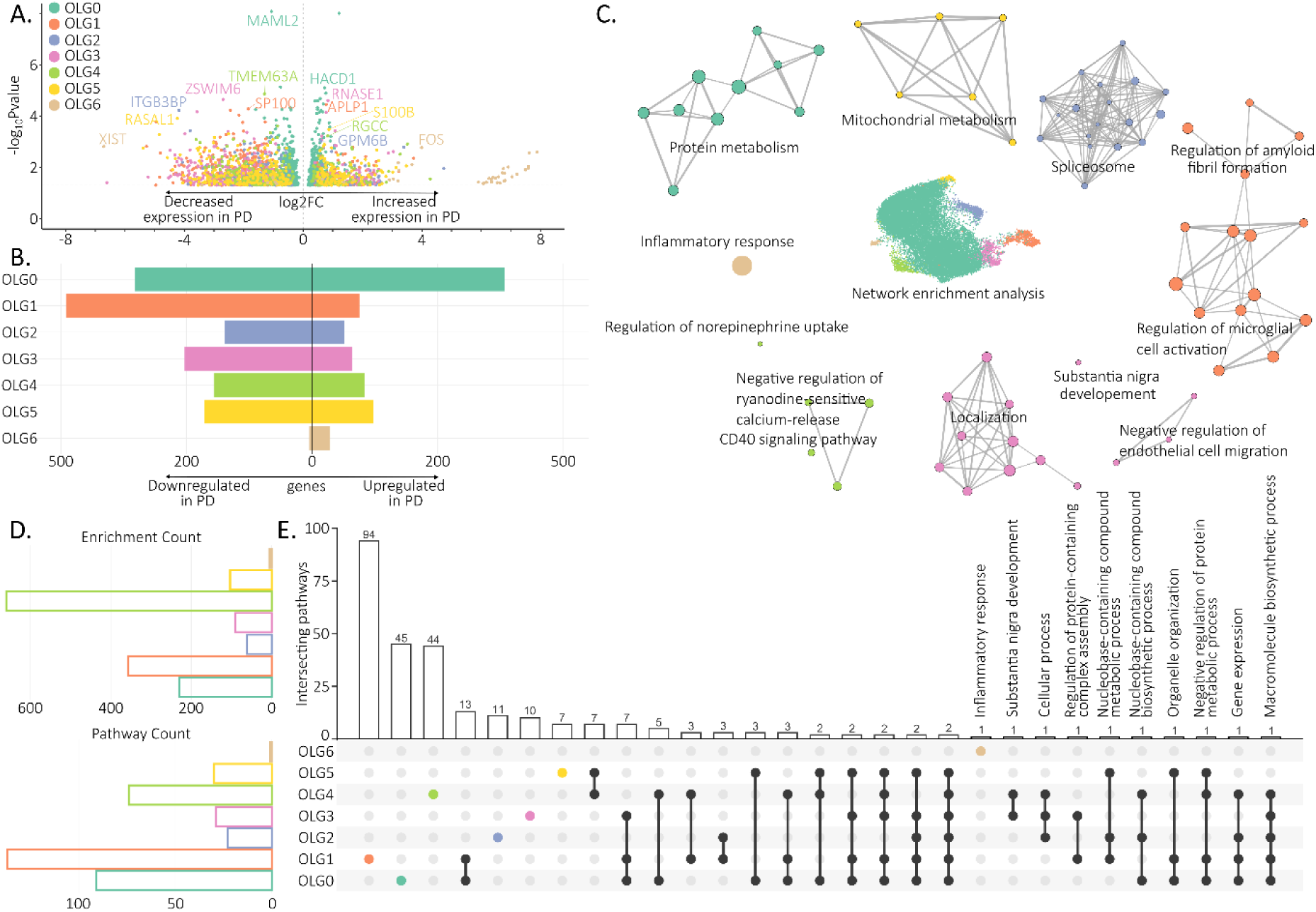
Transcriptomic analysis of oligodendrocyte subtypes. (A) Volcano plot of differentially expressed genes (DEG) in oligodendrocyte subtypes comparing PD and non-PD. Top upregulated and downregulated genes are labelled. Significant genes are coloured by subtype. (B) Bar plot showing counts of upregulated and downregulated DEGs, coloured by oligodendrocyte subtype. (C) Network representations of dysregulated GOBP pathways per subtype. Reduced and coloured by subtype. (D) Upper panel: bar plot of summed enrichment fold across all dysregulated pathways, coloured by subtype. Lower panel: bar plot of pathway counts, coloured by subtype. (E) Upset plot showing intersections of dysregulated pathways across subtypes. Intersections with *n* = 1 are labelled. PD Parkinson’s disease; OLG0-6 oligodendrocyte subtype 0-6.

To explore the functional relevance of these changes, we conducted gene ontology enrichment analysis on biological processes terms on the significant DEGs. In subtype 0, the most enriched pathways were related to protein metabolism, particularly protein folding and refolding (OLG0; Fig. S4 B,C). This fits with the interpretation of subtype 0 as a resting population, where PD-related stress appears to disrupt basic metabolic processes.

The distinction between the two observed inflammatory subtypes becomes clearer at the pathway level. In subtype inflammatory I, we observe enrichment for terms related to regulation of microglial cell-mediated cytotoxicity and microglial cell activation, with more specific pathways pointing to the negative regulation of amyloid formation and clearance processes (OLG1; Fig. S4 E,F). This aligns with previous studies reporting the aggregation of both α-synuclein and β-amyloid oligomers in oligodendrocytes, sometimes even more than in neurons [1, 29]. In contrast, subtype inflammatory II shows enrichment for pathways related to cell migration (OLG3; Fig. S4 K,L). This suggests that while subtype inflammatory I may be involved in the inflammatory clearance of protein aggregates, subtype inflammatory II might participate in inflammatory cell recruitment or migration, potentially across the blood-brain barrier.

In oligodendrocyte proliferative subtype, PD-related enrichment is observed in pathways linked to mRNA processing and RNA splicing (OLG2; Fig. S4 H,I). This supports a shift toward heightened transcriptional regulation, possibly reflecting increased demands for transcriptome reprogramming during pathological activation or stress responses [30, 31].

In the synaptic subtype, in PD, we show an upregulation in several relevant pathways (OLG4; Fig. S4 N,O). Enhanced regulation of norepinephrine uptake suggests a potential role in modulating axonal activity or stress signalling and it is also linked to calcium-metabolism [32]. Following, enrichment in calcium-release-related pathways points to altered intracellular signalling dynamics, which are tightly coupled to both dysregulated synaptic activity and oligodendrocyte support of axons in PD [33]. CD40 signalling, typically linked to immune responses in neurodegenerative diseases [33], may reflect inflammatory cross-talk with surrounding glial cells. Altered regulation of polyamine transport, involved in cell growth and repair, could indicate stress adaptation or disrupted resting, and might be further indication of the aforementioned subtype diversity continuum.

In the myelinating subtype, we find PD-related enrichment in pathways pointing to mitochondrial metabolic dysfunction (OLG5; Fig. S4 Q,R). Given the high energetic cost of myelin production and maintenance, even mild mitochondrial impairment can compromise oligodendrocyte function [34, 35]. This may contribute to the progressive loss of myelin integrity often reported in PD cortical regions [36].

When we examined the total number of enriched pathways across all subtypes, subtype inflammatory I displayed the highest count (Fig. 4 D). This aligns with its inflammatory profile, particularly its involvement in microglial cell-mediated cytotoxicity and responses to fibrillar deposit clearance. However, when considering the overall enrichment strength indicating the most pronounced dysregulation regardless of the number of pathways, the synaptic subtype ranked highest (Fig. 4 D). This synaptic-associated subtype exhibited marked disruption in pathways related to oligodendrocyte synaptic support in PD. At the same time, there was little overlap between the enriched pathways, suggesting that each human oligodendrocyte subtype, unlike mouse oligodendrocytes, engages in distinct molecular responses in PD, reflecting their functional specializations and divergent roles in disease pathology.

Overall, PD induces distinct functional disruptions across oligodendrocyte subtypes, reflecting diverse roles in stress response, inflammation, and metabolic vulnerability (Fig. 4 C). These findings highlight subtype-specific contributions to disease pathology.

### Disrupted cell signalling networks across oligodendrocyte subtypes and major neural cell types in PD

We next examined how dysregulated gene expression may alter intercellular communication, inferring signalling networks with CellChat [37]. It allows the identification of both outgoing and incoming communication patterns across multiple cell types and states. To this end, we used the oligodendrocyte subtypes alongside the 11 other cell populations present in the human dataset (Fig. 2 A).

A global view of signalling interactions revealed a noticeable reduction in both number and strength of intercellular communications in PD compared to non-PD (Fig. 5 A). In PD, both the number of interactions and their overall strength were reduced, aligning with our differential expression results that showed a predominance of downregulated genes across most oligodendrocyte subtypes. Among the pathways most affected were six major signalling routes. The APP (amyloid precursor protein) pathway, implicated in axon growth and synaptic plasticity, plays a role in neurodegenerative processes when dysregulated [38]. The CNTN (Contactin) pathway is critical for cell adhesion and myelination, particularly at axon-glia junctions [39, 40]. THBS (Thrombospondin) signalling is involved in synapse formation and extracellular matrix remodelling, often responding to injury or stress [41]. PECAM1 (Platelet Endothelial Cell Adhesion Molecule) is linked to immune cell transmigration across the blood–brain barrier and vascular inflammation [42]. The NEGR (neuronal growth regulator) pathway supports neurite outgrowth and synaptic stability, especially during development and repair [43]. Lastly, BAFF (B cell-activating factor) is associated with immune modulation and glial responses in neuroinflammation [44]. Notably, BAFF signalling was entirely absent in the PD group (Fig. 5 B), and all six pathways showed reduced information flow in PD compared to non-PD samples (Fig. 5 C).

**Figure 5.**
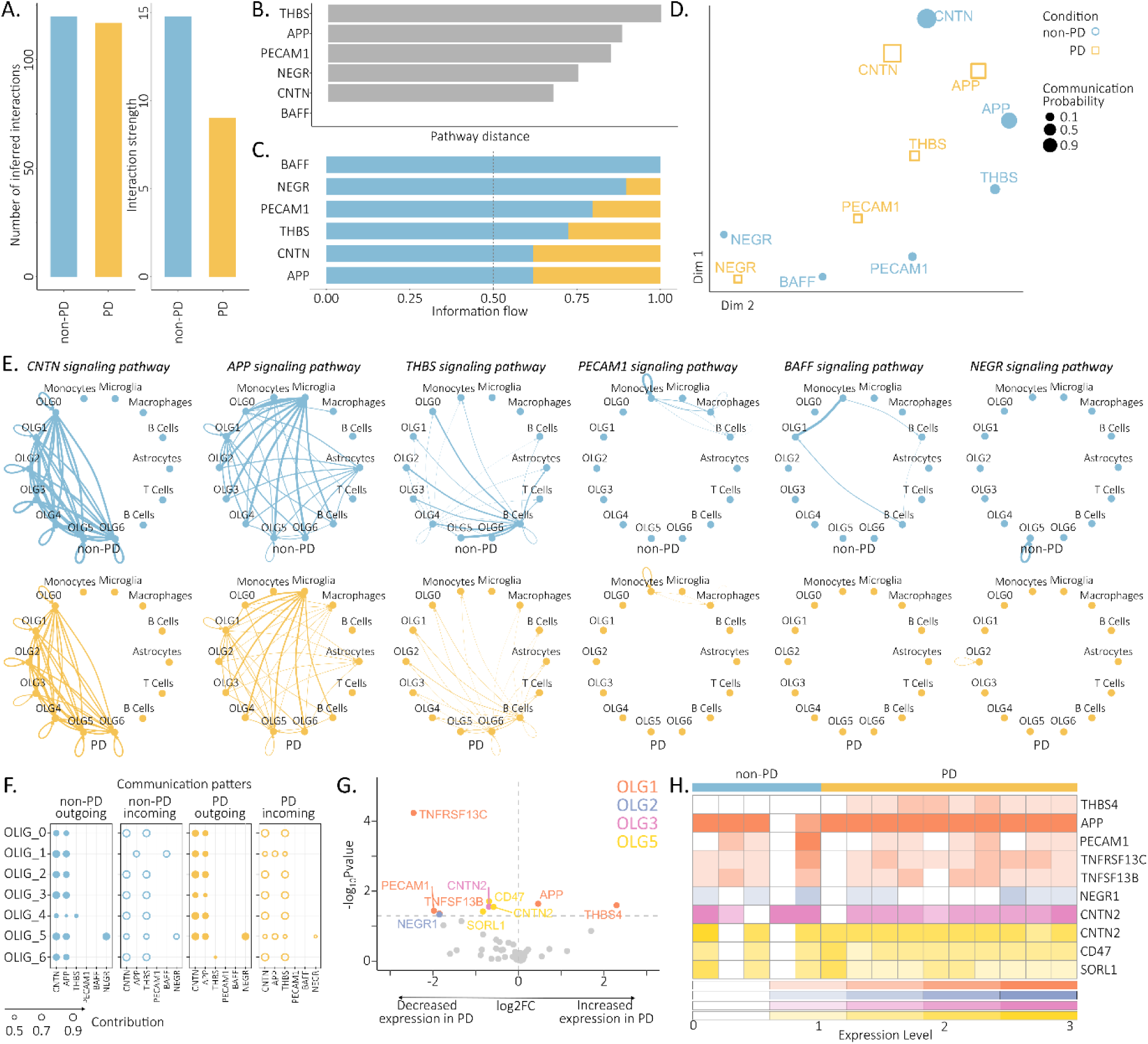
Comparative analysis of intercellular communication networks in PD vs non-PD. (A) Bar plot of the total number and overall strength of inferred cell–cell interactions in PD (yellow) vs non-PD (blue). (B) Bar plot of pathway distance in a shared two-dimensional space, showing whether equivalent signalling routes in PD and non-PD display similar patterns. (C) Bar plot of altered signalling pathways, comparing information flow, defined as the sum of communication probabilities across all cell group pairs, for each pathway; PD (yellow); non-PD (blue). (D) Signalling networks with the largest differences in functional similarity, based on Euclidean distance between shared pathways / cell-types. (E) Circle plots of the topology of each signalling pathway in PD (yellow) and non-PD (blue). (F) Summary for oligodendrocyte subtypes of overall communication patterns in PD (yellow) and non-PD (blue). (G) Volcano plot of differentially expressed genes (DEGs) enriched in signalling pathways. Significant genes are coloured by cluster. (H) Heatmap of normalized patient-level expression values for DEGs enriched in signalling pathways, with rows coloured by cluster. PD Parkinson’s disease; OLG0-6 oligodendrocyte subtype 0-6.

Additionally, we compared signalling activity between PD and non-PD and found that CNTN and APP pathways showed the strongest predicted interactions (Fig. 5 D). This is particularly important as APP signalling has been linked to PD mechanisms through its interaction with LRRK2, which modulates APP phosphorylation and enhances transcriptional activity of its intracellular domain, a process associated with dopaminergic neuronal loss under pathogenic conditions [45]. In parallel, disruption of the CNTN pathway reflects compromised axon–myelin integrity, as reduced levels of CNTN2 and associated nodal proteins impair neuronal communication during demyelination [46]. Together, these alterations point to convergent effects on neurodegenerative progression.

Breaking down these pathways across cell types, CNTN, APP, and THBS signalling were primarily involved in both inter- and intra-oligodendrocyte subtype communication, while PECAM1, NEGR, and BAFF were mostly restricted to inter-cellular interactions between oligodendrocytes and other brain cell types (Fig. 5 E). PD samples showed both fewer connections and weaker interaction strengths compared to non-PD (Fig. 5 F), consistent with the broader reduction in transcriptomic and signalling activity observed in the progressed disease context (Fig. 5 A).

After identifying which DEGs encode ligands or receptors involved in signalling pathways (Fig. S5 A,B), we examined their expression across oligodendrocyte subtypes (Fig. 5 G). In subtype inflammatory I, we observed upregulation of APP and THBS4, and downregulation of PECAM1, TNFRSF13B and TNFRSF13C (Fig. 5 G,H). This points to a selective dysregulation of APP, PECAM1, THBS, and BAFF signalling pathways in this subtype. The APP pathway implicated in axon growth and synaptic plasticity, is well-known for its central role in Alzheimer’s disease and neuroinflammation [47], and our findings suggest it also contributes to PD pathology. In line with recent studies [42, 44], our results highlight a functional role for BAFF and PECAM1 not only in immune modulation and glial activation during neuroinflammation, but also in mediating disrupted cell-cell communication in PD. In subtype inflammatory II, CNTN2 was downregulated (Fig. 5 G,H). As CNTN2 is involved in axon-glia adhesion and myelination [39, 40], this suggests that inflammatory oligodendrocytes may lose structural integrity and functional support, exacerbating neuronal vulnerability in PD.

In oligodendrocyte proinflammatory subtype, NEGR1 was downregulated (Fig. 5 G,H). As NEGR1 supports neurite growth and synaptic stability [43], its loss may hinder axonal repair and compromise synaptic integrity in PD. This is further seen in oligodendrocyte myelinating subtype, where we identified the downregulation of SORL1, CD47, and CNTN2, converging on the APP, CNTN, and THBS pathways (Fig. 5 G,H). As these pathways govern axon-glia contact, synapse formation, and neuroprotective signalling [38–41], their depletion in these subtypes suggests impaired structural support and compromised myelin maintenance contributing to the cortical atrophy observed in PD.

### Oligodendrocyte subtype transcriptomic shifts at late-stage mirror clinical severity and treatment load in PD

We next assessed whether oligodendrocyte-specific gene expression patterns in PD relate to clinical measures of disease progression in patients. To assess the disease state, we selected the following measurements: disease duration, Hoehn and Yahr stage, sex, UPDRS scores (off, on, and response), and levodopa equivalent daily dose (LEDD). Disease duration reflects the temporal progression of neurodegeneration and clinical symptoms [48]. Clinical symptoms, including tremor, rigidity, bradykinesia, postural instability, and gait difficulties, can be assessed on the Hoehn and Yahr scale, a standardized measure of motor symptom severity, categorizing patients into stages that reflect the disease progression [49]. Sex is a known factor to influence disease manifestation, progression and the severity of complications [50]. UPDRS on and off medication and its difference assess motor function and treatment response, respectively [51]. LEDD was developed as a standardized measure to quantify total dopaminergic medication intake, facilitating comparisons across diverse treatment regimens and helping to assess therapy impact despite variability in medication types and dosages [52]. We computed module eigengenes, summary measures of dysregulated gene expression [53] within each oligodendrocyte subtype, and evaluated their correlation with clinical metrics at the individual patient level (Fig. 6 A).

**Figure 6.**
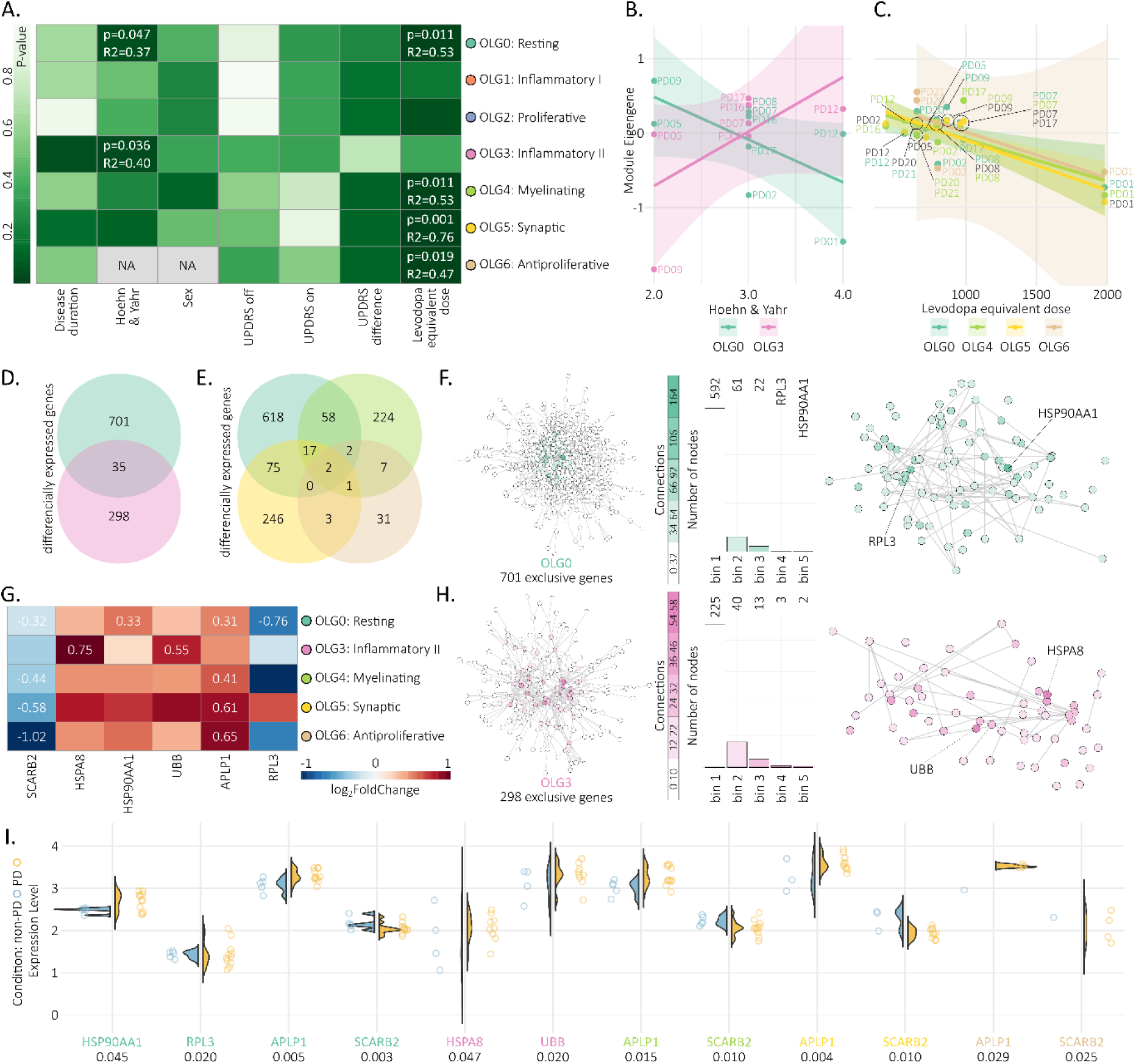
Clinical relevance of oligodendrocyte subtypes transcriptomic changes. (A) Heatmap showing patient-wise Pearson correlation between module eigengenes (WGCNA) and clinical measures, coloured by p-value. Labelled by oligodendrocyte subtype. (B) Scatter plot of significant correlations with Hoehn & Yahr stage; coloured by cell type. (C) Scatter plot of significant correlations with levodopa equivalent dose; coloured by cell type; oligodendrocyte subtype 5 shown in yellow and black. (D) Venn diagram of differentially expressed genes (DEGs) driving the correlation in (b), for subtypes 0 and 3. (E) Venn diagram of DEGs driving the correlation in (C), for subtypes 0, 4, 5, and 6. (F) Protein–protein interaction network of DEGs in subtype 0 inversely correlated with Hoehn and Yahr. Network coloured by connectivity. Bar plot shows node counts. Second network showing only nodes in bins 2–5. Top two hub genes labelled. (G) Heatmap of expression levels for six selected genes. Coloured by differential expression level in PD compared to non-PD. (H) Protein–protein interaction network of DEGs in subtype 3 directly correlated with Hoehn & Yahr, visualized as in (F). (I) Violin plots showing patient-averaged expression level of six selected genes across cohorts. PD Parkinson’s disease. OLG0-6 oligodendrocyte subtype 0-6.

We identified two significant correlations with the Hoehn and Yahr scale (Fig. 6 B): a negative correlation with the resting subtype, and a positive correlation with inflammatory II subtype; suggesting that as disease severity increases, homeostatic functions decline facilitating inflammatory responses in oligodendrocytes. This reflects a shift in cellular states linked to PD progression towards a late-stage. To delve deeper into these correlations, we examined the DEGs characteristic of these subtypes and assessed their overlap (Fig. 6 D). We hypothesized that genes exclusively dysregulated in the resting subtype contribute to the negative correlation with disease severity, while those unique to inflammatory II subtype drive the positive correlation (Fig. 5 D). In the resting subtype, we identified 701 exclusive DEGs, which were further analysed for protein-protein interactions to elucidate their functional roles (Fig. 6 F) [54]. These DEGs were assigned into five connectivity-based bins, with the top two hubs identified as HSP90AA1 and RPL3. HSP90AA1 encodes a molecular chaperone involved in protein folding and stress responses critical for cellular homeostasis, while RPL3 encodes a ribosomal protein essential for protein synthesis and cellular metabolism [16]. Conversely, we identified 298 exclusive DEGs in inflammatory II subtype, which we similarly analysed for protein-protein interactions (Fig. 6 H). The top hubs in this network were HSPA8 and UBB. HSPA8 encodes a heat shock protein that plays a key role in protein quality control and stress response, and is reported to increase monomeric α-synuclein levels [55]. UBB encodes ubiquitin B, which is crucial for tagging damaged or misfolded proteins for degradation, thereby maintaining proteostasis under pathological conditions and is notably also linked to neurodegenerative diseases, especially PD [56, 57].

Conversely, we identified four significant positive correlations between LEDD and oligodendrocyte subtypes: resting, synaptic, myelinating and anti-proliferative (Fig. 6 C), indicating that higher medication requirements coincide with more pronounced transcriptomic alterations. To dissect these associations, we examined the DEGs characteristic of these subtypes and assessed their overlap (Fig. 6 E), revealing two shared genes of particular interest. SCARB2 encodes lysosomal integral membrane protein type 2, essential for lysosomal function and previously linked to PD risk via GBA1-related pathways [58]. Amyloid precursor-like protein 1 (APLP1) participates in pathological α-synuclein transmission, with significantly higher levels detected in PD patients [59].

We next evaluated the expression patterns of these six hub genes. Notably, only RPL3 and SCARB2 were downregulated (Fig. 6 G,I), while HSP90AA1, HSPA8, UBB, and APLP1 were upregulated. Collectively, the expression profile of these hub genes delineates a convergent disturbance across multiple proteostatic and metabolic axes in PD. Downregulation of RPL3 and SCARB2 points to impaired ribosomal protein synthesis and lysosomal degradation capacity, two pillars of cellular maintenance, potentially undermining oligodendrocyte resilience.

## DISCUSSION

In this study we sought to uncover the contribution of oligodendrocytes to PD progression by defining disease stage–specific transcriptional signatures. To this end, we analysed cortical single-cell transcriptomes from early- and late-stage parkinsonian mouse models and compared them with PD and non-PD patient datasets. In total, we profiled 55,647 mouse and human oligodendrocytes, which resolved into distinct subtypes encompassing resting, inflammatory, proliferative, synaptic, and myelinating states. Across species, early disease in mice was marked by immune and inflammatory activation, while advanced stages in both mice and humans converged on metabolic dysfunction and disrupted proteostasis, including impaired ribosomal output, chaperone stress responses, ubiquitination deficits, and lysosomal perturbation. Patient samples presented a shift in cellular composition, with depletion of resting oligodendrocytes and expansion of inflammatory and proliferative states. Trajectory analysis supported a continuum of dynamic state transitions rather than discrete endpoints. These changes were accompanied by a global reduction in intercellular signalling. Importantly, clinical correlation analyses linked these oligodendrocyte state shifts to disease severity and treatment demand, highlighting their relevance to late-stage PD progression.

Defining oligodendrocyte subtypes remains challenging due to the inherent plasticity of glial cells [25]. Unlike neurons, many glia cells retain the capacity to transition between states, which complicates the distinction between cell type and state [25, 60, 61]. Conceptually, these populations can be considered as subtypes, within which individual states represent dynamic positions [62]. Mouse samples showed a more homogeneous population distribution, while human samples revealed multiple mature oligodendrocyte subtypes, with a limited number of subtype-specific markers [25]. Instead, functional heterogeneity is reflected in patients with PD in the differential enrichment of pathways, underscoring the complexity and increased heterogeneity of oligodendrocyte biology in the merging disease context. The observed shifts affect key functions, including protein metabolism, synaptic support and mitochondrial integrity, inflammation and myelination.

Oligodendrocytes are increasingly recognized as active contributors to immune regulation, particularly in initiating inflammatory responses [63, 64]. They express diverse immunoregulatory factors, including cytokines and chemokines, positioning them as active participants in neuroinflammation [63, 65].

Consistent with this, our analysis supports this concept with both mouse and human data. In the early-stage of the disease we reported in our mouse model a strong upregulation of genes and pathways related to inflammatory activation. This activity is not recorded at the late-stage of the disease. Conversely in our patients, oligodendrocyte inflammatory I subtype displayed strong dysregulation of pathways associated with α-synuclein aggregation as well as altered crosstalk with microglia, aligning with the idea of oligodendrocytes as key responders to extracellular pathogenic signals. Furthermore, oligodendrocyte inflammatory II subtype was enriched for pathways linked to mRNA processing, intracellular localization, and transmission, suggesting an intrinsic contribution of oligodendrocytes to α-synuclein accumulation. This would suggest that on one side that the pathology progresses faster and rather directly in mice. On the other side, in humans we encounter a more heterogenous pattern due to higher complexity of the system and compensatory potential.

In PD, one proposed mechanism underlying oligodendrocyte pathology is the transmission of misfolded α-synuclein from neurons, which could explain the prominent inclusions observed in these cells [66]. In vitro evidence shows that rodent oligodendrocyte lines can internalize both monomeric and oligomeric α-synuclein [67]. Similarly, abundant oligodendrocyte α-synuclein inclusions were reported in human PD tissue, initially attributed solely to uptake of neuronal protein [67, 68]. However, newer approaches demonstrate that oligodendrocytes also produce α-synuclein endogenously at the mRNA and protein levels [69].

In the context of spreading, α-synuclein dynamics appear closely linked to oligodendrocyte dysfunction and disease progression. In our data, APLP1 was consistently upregulated and correlated with clinical markers of disease severity, supporting its role in facilitating α-synuclein propagation within oligodendrocytes. At the same time, disruption of the APP pathway, involved in axonal growth and synaptic plasticity, points to broader alterations in protein processing and intercellular signalling under pathological conditions. In parallel, attenuation of CNTN signalling, a pathway essential for axon–glia interactions and myelination, suggests a reduced capacity for structural repair, consistent with a loss of cellular resilience. These changes occur alongside downregulation of RPL3 and SCARB2, indicating impaired ribosomal function and lysosomal degradation, respectively, and further reinforcing the accumulation of proteotoxic stress as the disease advances. Notably, the conservation of RPL3-associated signatures across species highlights its potential as a robust marker of oligodendrocyte dysfunction in PD.

Transcriptomic changes in oligodendrocytes at late-stage PD increasingly align with clinical manifestations, linking cellular dysfunction to disease severity. Imaging studies show that reduced myelin integrity correlates with motor impairment (UPDRS) and widespread white matter alterations track with clinical progression [70, 71]. Additional evidence connects oligodendrocyte dysfunction to cognitive decline, neuropsychiatric symptoms, and disease outcomes [72, 73]. In line with this, our human data reveal that resting oligodendrocytes negatively correlate with Hoehn and Yahr stage, whereas inflammatory subtypes show the opposite trend, indicating a shift in cellular states with disease progression. Furthermore, multiple subtypes positively correlate with levodopa equivalent daily dose (LEDD), suggesting that increased therapeutic demand accompanies deeper transcriptional reprogramming.

Several limitations should be considered when interpreting these findings and their clinical relevance. Validation across additional PD mouse models with distinct pathogenic mechanisms is needed to confirm the robustness and generalizability of the identified oligodendrocyte states. The current focus on cortical, anatomically matched samples also restricts insight into region-specific and systemic disease effects, warranting extension to other central nervous system regions and peripheral tissues. Moreover, clinical correlations do not resolve whether oligodendrocyte reprogramming drives or results from disease progression. Addressing this will require longitudinal studies, in vivo functional perturbation of oligodendrocyte subtypes, and multimodal validation combining imaging, biofluid markers, and patient-derived models to establish causality and therapeutic relevance.

This study underscores the clinical and mechanistic relevance of oligodendrocytes in PD. Subtype-specific analyses revealed that declining support and salient inflammatory activity parallel disease severity across species, while broader transcriptional changes correlate with increasing treatment requirements. In line with previous evidence, oligodendrocyte subtypes exacerbate the underlying pathology in PD, through impaired metabolic support, dysregulated stress and immune signalling, and defective autophagy. APP and CNTN pathways sit at the signalling intersection, linking α-synuclein pathology with axon–glia communication and structural support, and thereby shaping disease progression. Together, these findings highlight oligodendrocyte dysfunction as a key translatable factor from mice to patients in PD progression and pathology.

## MATERIALS AND METHODS

### Study design

In this study we investigated cell type–specific transcriptomic alterations in PD. We compared single-cell RNA sequencing of human DLPFC tissue with a parkinsonian mouse model overexpressing human mutant A53T α-synuclein [15]. We hypothesized that oligodendrocytes display transcriptional dysregulation contributing to PD pathology progression. In total, 16 patients including 11 with PD and 5 non-PD were included (Table S1). In parallel, mouse experiments included stereotactic rAAV2/7 injections on 20 mice, with 22 controls. Euthanasia at early (4 weeks) or late (8 weeks) post viral injections was conducted to reveal pathological disease stages. Cortical tissue from 42 C57BL/6N mice was dissected and pooled to meet experimental input requirements. The analysis pipeline included capturing high-quality single-cell transcriptomes [19], oligodendrocyte subtype identification [19], pseudotime trajectories [26], differential gene expression [74], co-expression networks [53], cell–cell communication [37] and association analyses. Sample sizes were guided by prior similar studies in our group. No samples were excluded.

### Ethics

Written informed consent was obtained from all participants prior to enrolment. The study on patients was conducted in accordance with the Declaration of Helsinki and approved by the local ethics committee (837.208.17 (11042)). Participants received no compensation. All procedures involving the care and treatment of laboratory animals were approved by the local ethics committee (23177–07/G 19–1-081) and complied with the European and German national regulations (European Communities Council Directive, 86/609/EEC).

### Mice

Male C57BL/6 N mice (age: 2 to 4 months, 24–32 g) were obtained from the Translational Animal Research Center (TARC, Mainz, Germany) and housed in the Institute of Physiology, University Medical Center of the Johannes Gutenberg University Mainz (Mainz, Germany) facility. Mice were maintained at a 12-hour dark/light cycle and received standard food and water at libitum. In total, 42 animals were planned for this study. Due to meeting the minimum weight required for the single-cell pipeline, two mice were pooled for every single experimental workflow, resulting in 21 datapoints. No animal or sample was excluded from the study.

### Surgical Preparation and Euthanasia

For this study 20 mice were injected with human A53T α-synuclein rAAV2/7 vector to create a parkinsonian model as described elsewhere [15]. In brief, the animals were anesthetized with 1.5–2% isoflurane and secured in a stereotactic frame (Kopf Instruments). After shaving and disinfecting the scalp, a midline incision was made and the skull was cleaned with 70% ethanol. A burr hole was drilled over the left substantia nigra using stereotactic coordinates (anteroposterior −3.1 mm, mediolateral −1.2 mm relative to bregma, dorsoventral −4.0 mm from the dura) based on the mouse brain atlas [75]. Using a glass micropipette on a micromanipulator, 2 μl of viral vector dose 4,0E + 11 GC/ml were delivered at 0.25 μl/min [15]. The pipette was left in place for 15 minutes to prevent reflux before being slowly withdrawn. Mice were sacrificed after an observational period (of either 4 or 8 weeks) after the acute experiment, using an overdose of anaesthesia. Based on published results 4 weeks of viral overexpression is estimated as an early disease stage, while 8 weeks of viral overexpression is estimated as a late disease stage [15]. In parallel, 22 mice were observed for respective intervals of time as controls and sacrificed the same manner. Samples of dissected cortical tissue were combined from two mice to reach the recommended weight for the subsequent experimental workflow.

### Patient Selection and Description

In total, 16 patients (mean age: 56.86 ± 12.94 years; 9 female) were enrolled, including 11 with PD (mean age: 58.82 ± 13.86 years; mean disease duration: 9.73 ± 4.88 years) and 5 non-PD individuals (mean age: 57.8 ± 16.2 years; mean disease duration: 13.4 ± 5.6 years). Patients included in previous published work have been included (14 patients) [12, 27] in addition to newly recruited ones. All patients were recruited at the University Medical Center of the Johannes Gutenberg University Mainz. Clinical evaluations were conducted by a movement disorders specialist at the Department of Neurology. Prior to enrolment, all patients undergoing DBS received comprehensive screening and met established DBS eligibility criteria [76]. A clinical neuropsychologist excluded cognitive impairment (MoCA >24; Mattis DRS >138). PD diagnoses were confirmed according to the UK Parkinson’s Disease Society Brain Bank criteria. Non-PD individuals showed no evidence of major neurodegenerative or metabolic disorders and comprised a heterogeneous group including dystonia, essential tremor, and obsessive-compulsive disorder. Clinical characteristics of the donors have been elaborated in Table S1.

### Sample Processing

DLPFC tissue samples (50–100 mg) were collected from the patients intraoperatively by the neurosurgeon during DBS and immediately transferred into sterile 50 ml centrifuge tubes containing Hanks’ Balanced Salt Solution (HBSS). Mouse tissue samples (50–100 mg) were obtained following dissection after euthanasia, as described above, and handled similarly.

Samples were kept on ice and transported immediately to the laboratory, where they were washed three additional times in HBSS to remove blood. They were subsequently preserved in GEXSCOPE tissue preservation solution (Singleron Biotechnologies) at 4°C to await further processing. All samples were processed within 24 hours. In brief, tissues underwent mechanical and enzymatic dissociation (Tumor Dissociation Kit, 130-095-929, Miltenyi), followed by sequential removal of debris (Debris Removal Solution, 130-109-398 Miltenyi), red blood cells (Red Blood Cell Lysis Solution, 130-094-183, Miltenyi), and dead cells (Dead Cell Removal Kit, 130-090-101, Miltenyi). Single-cell suspensions were quantified using automated cell counting (LUNA-FL Dual Fluorescence Cell Counter) with viability staining (Acridine orange stain, F23002). Because it relies on combined mechanical and enzymatic dissociation, the microwell SCOPE-chip platform (Singleron Biotechnologies) is less suitable for human neurons, which are relatively large and particularly susceptible to membrane damage.

Single-cell gene expression libraries were prepared following manufacurer’s protocols for cell partitioning, lysis, reverse transcription, and cDNA amplification (Singleron Biotechnologies). Individual cells were loaded onto a SCOPE-chip partitioning single cells into microwells, where they were lysed and mRNA was captured by barcoded beads with transcript-specific unique molecular identifiers (UMIs). Amplified cDNA was enzymatically fragmented, end-repaired, and ligated to adapters with 3′ A-overhangs. P5 and P7 sequences and sample indices were added during final PCR amplification. Fragment size was assessed with the Bioanalyzer High-Sensitivity DNA Kit (Agilent, CA, USA), and library concentrations were measured using the Qubit dsDNA HS Assay Kit (Thermo Fisher Scientific, MA, USA). All samples met quality control criteria and yielded sufficient material for sequencing. Final single-cell RNA-sequencing libraries were outsourced for paired-end sequencing on a NovaSeq 6000 platform.

### Data Processing

Demultiplexed raw reads were processed using CeleScope (v1.8.1, Singleron Biotechnologies) with the GRCh38 human reference genome or the GRCm39 Mus musculus reference for alignment, quality filtering, barcode assignment, and UMI quantification.

In brief, cells expressing at least 200 unique molecular identifiers in a minimum of three cells had sufficient complexity and were retained for downstream analysis. Gene expression data were normalized using the NormalizeData() function in Seurat v4.1.0 [19]. Quality control included two main steps: (1) identification and removal of doublets and multiplets with DoubletFinder v2.0.4 on a per-sample basis [77], and (2) exclusion of cells with mitochondrial content >10% and fewer than 500 detected genes [19]. DoubletFinder parameters for generating artificial doublets and classifying true doublets followed default recommendations. After further filtering, high-quality single cells were retained. To correct for batch effects across samples, data were integrated using Seurat’s FindIntegrationAnchors() and IntegrateData() functions, based on Canonical Correlation Analysis (CCA) with default settings (2,000 features; LogNormalize; scale = TRUE; reduction = “cca”; dims = 1:30) [78]. This ensured appropriate integration according to molecular characteristics and not by sample and or batch.

Standard downstream analysis included PCA, graph-based clustering, and UMAP dimensionality reduction. Resolution was refined iteratively until distinct cell populations were resolved. We used a resolution of 0.7 for the human and 0.5 for the mouse dataset. Clusters were annotated manually based on established marker genes and curated databases [20] as well as our previous experience [12, 27], and consolidated major cell types.

Cells labelled as oligodendrocytes were subset and reanalysed, including PCA, graph-based clustering, and UMAP dimensionality reduction, performed at a resolution of 0.1, which resolved transcriptionally distinct subtypes. Cluster-specific markers were identified using FindMarkers() function from the Seurat “RNA” assay using log-normalized counts and the default Wilcoxon rank-sum test in Seurat v4.1.0 (adjusted P < 0.05). Marker gene overlaps between clusters were assessed using UpSet plots (UpSetR v1.4.0) [79] and Euler diagrams (eulerr v7.0.2). Gene ontology enrichment for biological processes on marker genes was performed with PANTHER [80], and clusters were annotated accordingly.

Cell state trajectories for human oligodendrocytes were reconstructed using Monocle3 (v1.4.26) to infer dynamic transcriptional changes and potential lineage relationships among oligodendrocyte subtypes [26]. Briefly, cells were ordered in pseudotime based on high-variance genes using reversed graph embedding, allowing the identification of branch points that may correspond to distinct functional or developmental transitions. Genes that varied significantly along the pseudotime axis were further analysed using the expression values transformed as log1p(x).

Differential gene expression between PD and non-PD was analysed using a pseudobulk strategy followed by DESeq2 (v1.34.0) [74]. Enriched biological processes among differentially expressed genes were identified with PANTHER [80], visualized using UpSet plots for pathway overlaps [79], and enrichplot (v1.14.2) through clusterProfiler (v4.2.2) for network analysis [81].

Cell–cell communication for the human dataset was analysed using CellChat (v2.1.2), which integrates ligand–receptor expression data to infer signalling interactions between cell types [37]. Signalling probabilities were calculated for each oligodendrocyte subtype to identify key pathways and interaction changes between PD and non-PD. Results were visualized using network and circle plots to compare communication patterns across conditions. Key genes were explored based on their expression.

Weighted gene co-expression networks (WGCNA v1.72.1) were built for each human oligodendrocyte subtype using differentially expressed genes, applying an unsigned adjacency matrix with soft-thresholding and hierarchical clustering [53]. Module eigengenes were correlated with PD-relevant clinical traits. In modules showing significant associations, overlapping genes were identified via Venn diagrams (VennDiagram v1.7.3) [82] and queried in STRINGdb database for potential protein–protein interactions [54].

### Statistical Analysis

All statistical analyses were performed in RStudio (v1.4.1717) using established packages and pipelines, including Seurat, DESeq2, WGCNA, PANTHER, enrichR, CellChat, and STRINGdb. Seurat workflow was used for log-normalization to scale and stabilize variance, followed by graph-based Louvain clustering on PCA-reduced data [19]. Differential gene expression was assessed with DESeq2 using a negative binomial model, and p-values were adjusted with false discovery rate (FDR) correction [74]. Overrepresentation analyses were conducted using PANTHER and enrichR, applying Fisher’s exact test with FDR correction for multiple comparisons [80, 83]. Network and enrichment analyses, including STRINGdb [54], were performed depending on the specific research questions, ensuring robust and comprehensive statistical evaluation. WGCNA was used to construct gene co-expression networks by calculating pairwise gene correlations and identifying modules through hierarchical clustering [53]. Module–trait associations were tested using Pearson correlation with phenotypic data. Frequency differences of cell types were analyzed using linear mixed-effects models, which allowed assessment of different parameters while accounting for intra-group correlations. Bonferroni correction was applied to adjust for multiple comparisons.

## Author contributions

Conceptualization: GGE, SG

Methodology: HJL, TB, GGE, SG

Investigation: DM, MBG, SLK

Resources: DM, SK, MK

Data curation: DM, MBG, SLK, VA, JB

Visualization: DM

Supervision: HJL, GGE, SG

Writing – original draft: DM

Writing – review & editing: NS, YD, MBG, SLK, HJL, VA, JB, SK, MK, TB, MH, PLDJ, GGE, SG

Funding acquisition: SG

## Data Availability statement

Raw data generated in this work and any additional information required to re-analyze the data reported are available upon request from the lead contact.

## Code availability statement

All analyses were performed using openly available software and toolboxes and are available upon request from the lead contact.

