## Supplement for "Oligodendrocyte subtype diversity underlines clinical progression in Parkinson’s disease"

**This document includes:**

Supplementary figures 1-5

Supplementary table 1

### Supplemental Figures

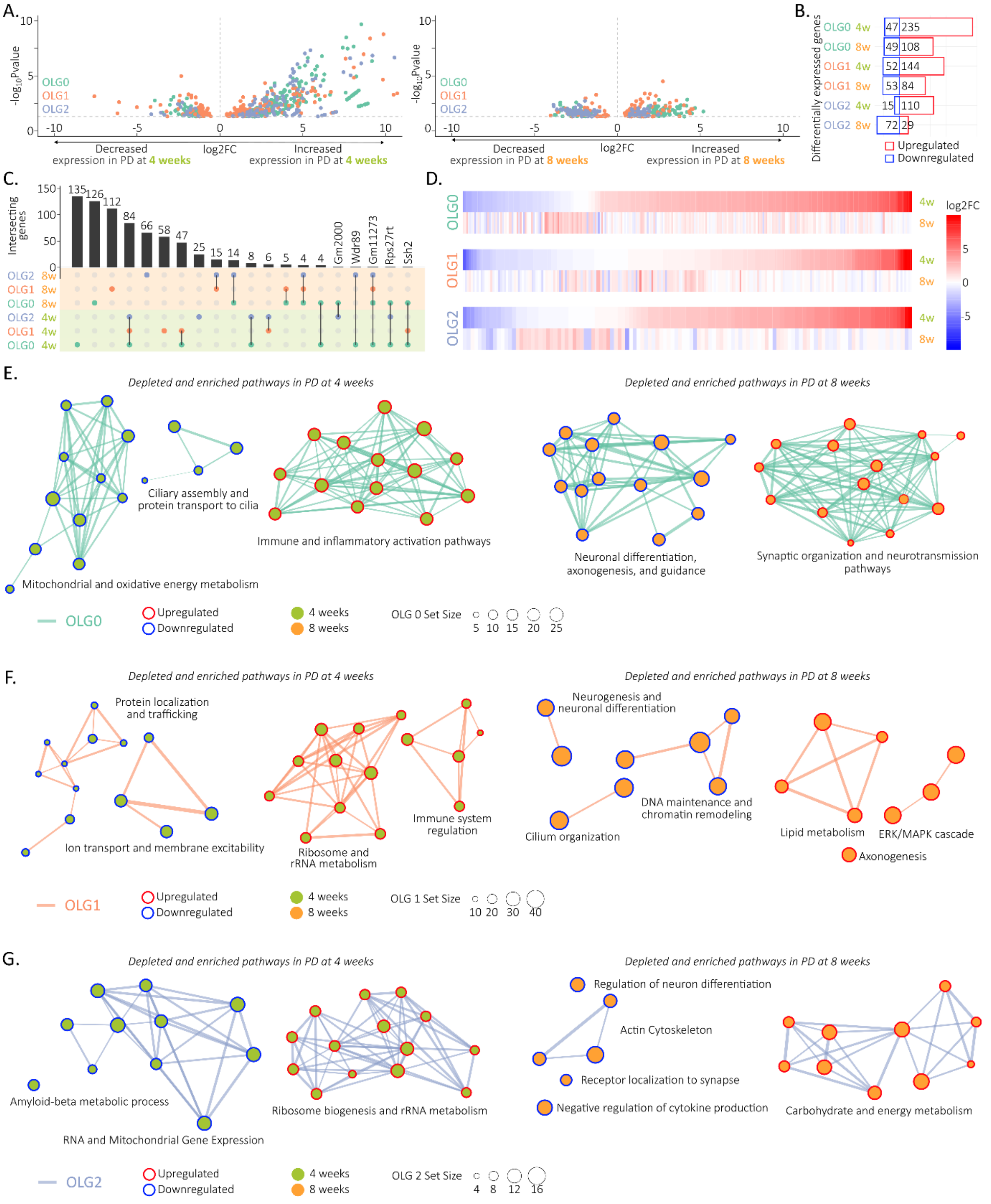


**Figure 1 Supplemental Legend. Differential gene expression across mouse oligodendrocyte subtypes**

(A) Volcano plot of differentially expressed genes (DEG) in mouse oligodendrocyte subtypes comparing PD (parkinsonian mouse model) and non-PD (control mice) at four weeks (left panel) and eight weeks (right panel) of observation. Significant genes are coloured by subtype. (B) Bar plot showing counts of upregulated and downregulated DEGs, grouped by mouse oligodendrocyte subtype and period of observation. (C) Upset plot displays intersections of DEGs across subtypes. (D) Heatmap displays expression differences of DEGs across subtypes. (E, F, G) Network representations of dysregulated Gene Ontology Biological Process pathways; grouped by subtype; colour of the connecting lines corresponds to the subtype; size of the dot corresponds to the set size of the pathway; colour of the dot corresponds to the period of observation; colour of the circle corresponds to the dysregulation (enriched or depleted). PD parkinsonian mouse model; non-PD control mice; OLG0-2 oligodendrocyte subtype 0-2.

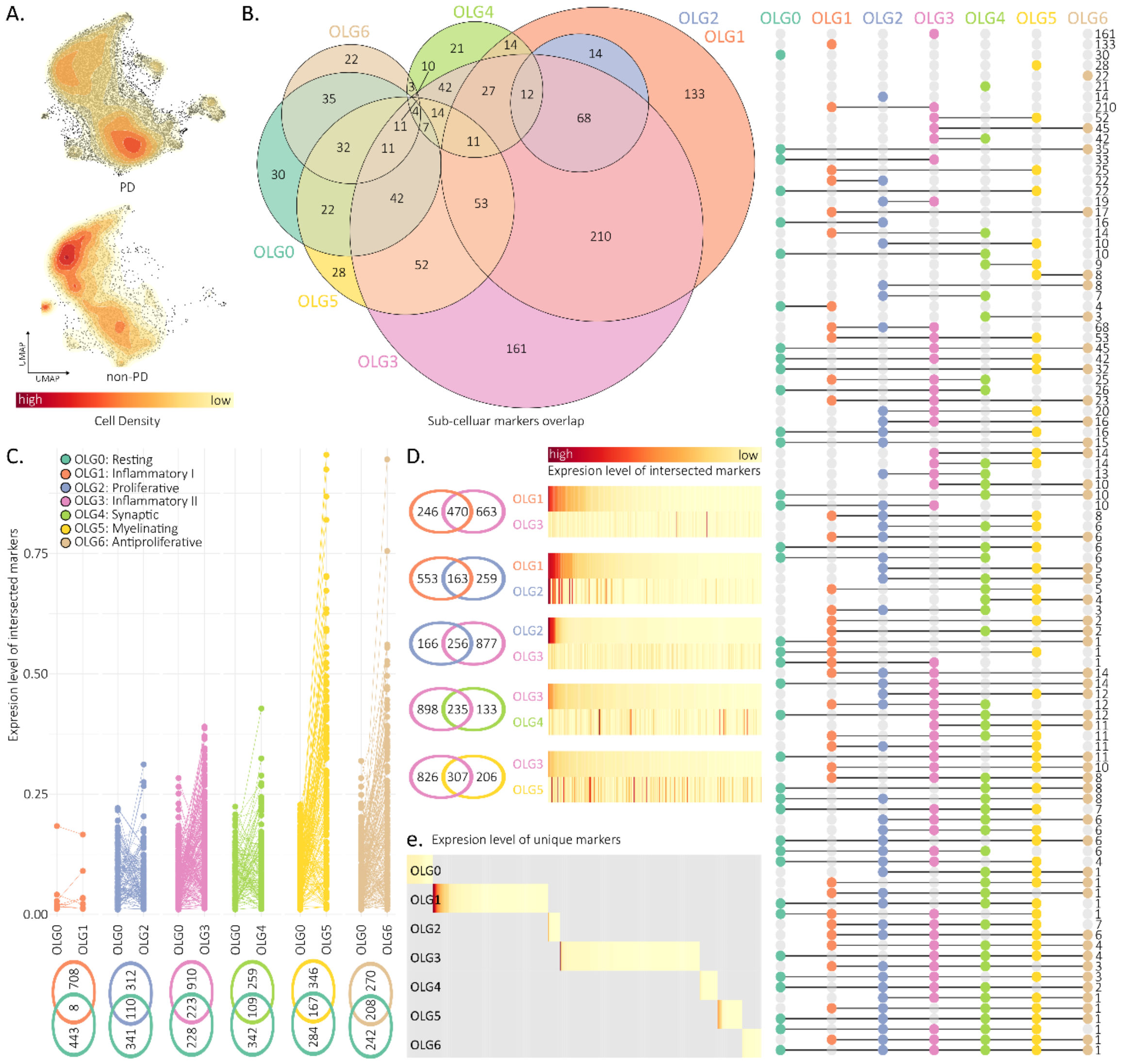


### Figure 2 Supplemental Legend. Marker genes expression across oligodendrocyte subtypes

(A) Density plot of oligodendrocyte cells by cohort, based on UMAP embedding. (B) Euler diagram of gene markers identified in each subtype, with circles coloured by subtype. Upset plot displays intersections of marker genes across subtypes. (C) Venn diagram of gene markers shared between subtype 0 and other oligodendrocyte subtypes. Line-connected dot plot shows expression differences of the shared markers across subtypes. (D) Venn diagram of gene markers shared among inflammatory oligodendrocyte subtypes. Heatmap displays expression differences of shared markers across these subtypes. (E) Heatmap of subtype-specific marker gene expression, of unique genes markers in each subtype. PD Parkinson's disease; UMAP Uniform Manifold Approximation and Projection; OLG0-6 oligodendrocyte subtype 0-6.

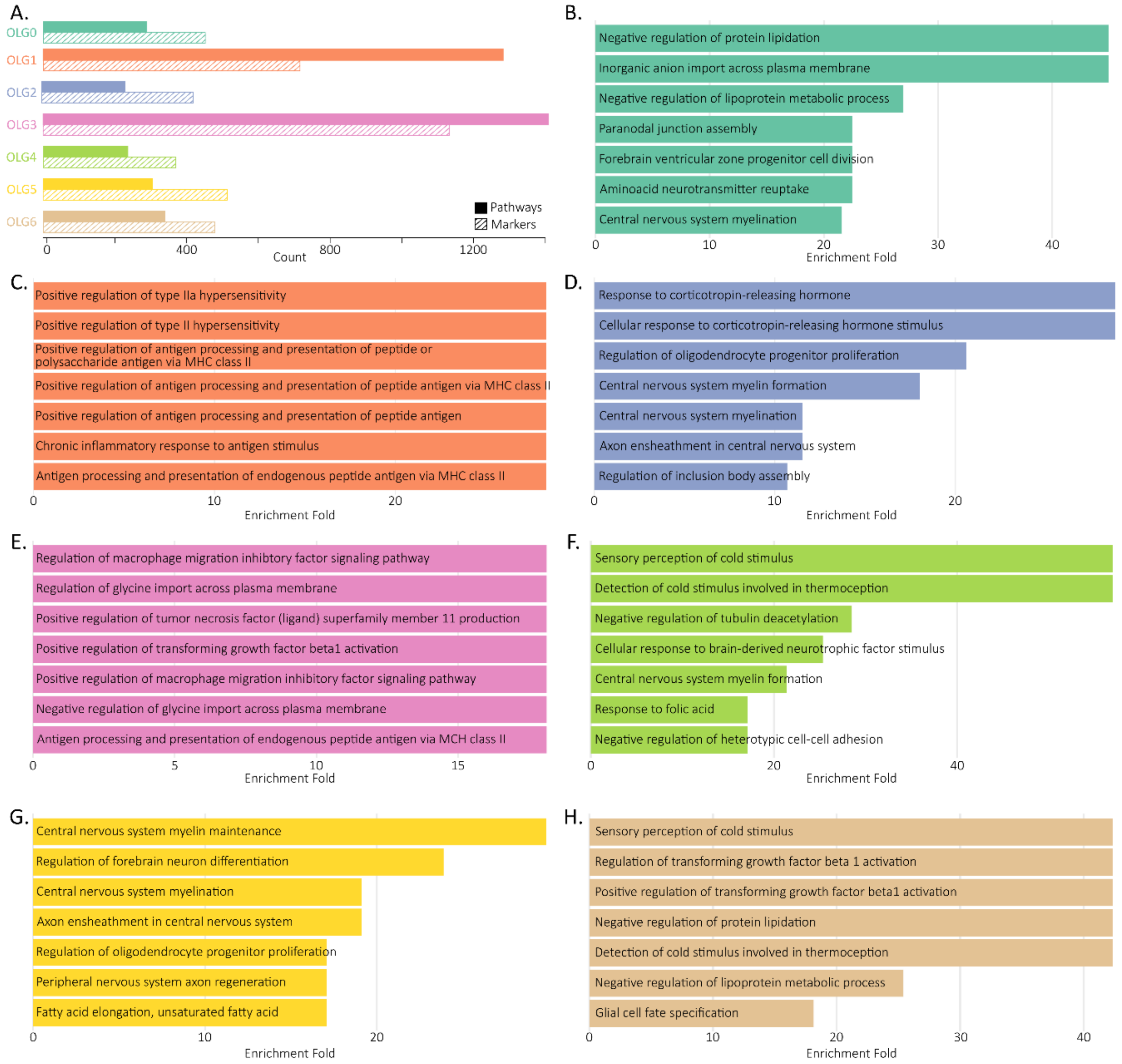


**Figure 3 Supplemental Legend. Gene ontology enrichment across marker genes of oligodendrocyte subtypes.**

(A) Bar plot showing the number of marker genes per oligodendrocyte subtype and the count of enriched Gene Ontology Biological Process (GOBP) pathways identified in each. (B–G) Top seven enriched GOBP pathways for each oligodendrocyte subtype (0–6), ranked in descending order of enrichment fold. Coloured by oligodendrocyte subtype. OLG0-6 oligodendrocyte subtype 0-6.

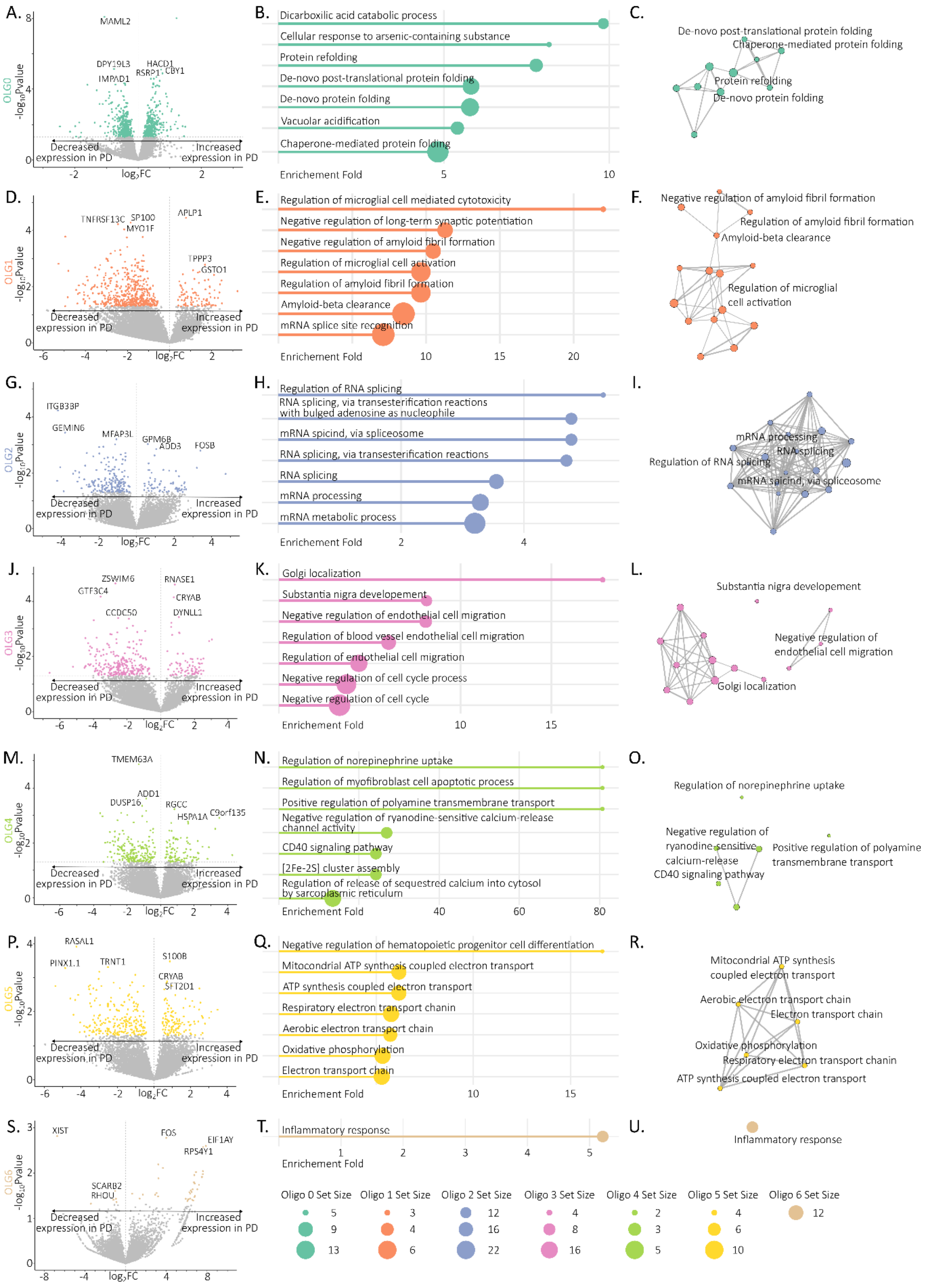


**Figure 4 Supplemental Legend. Differential gene expression and pathway enrichment across oligodendrocyte subtypes.**

(A, D, G, J, M, P, S) Volcano plots of differentially expressed genes (DEG) in oligodendrocyte subtypes 0–6 comparing PD and non-PD. Top three upregulated and downregulated genes are labelled. Significant genes are coloured according to subtype. (B, E, H, K, N, Q, T) Bar plots showing the top seven dysregulated Gene Ontology Biological Process (GOBP) pathways in each subtype, ranked by enrichment fold. Coloured by subtype. (C, F, I, L, O, R, U) Network representations of dysregulated GOBP pathways in each subtype. Coloured by subtype.

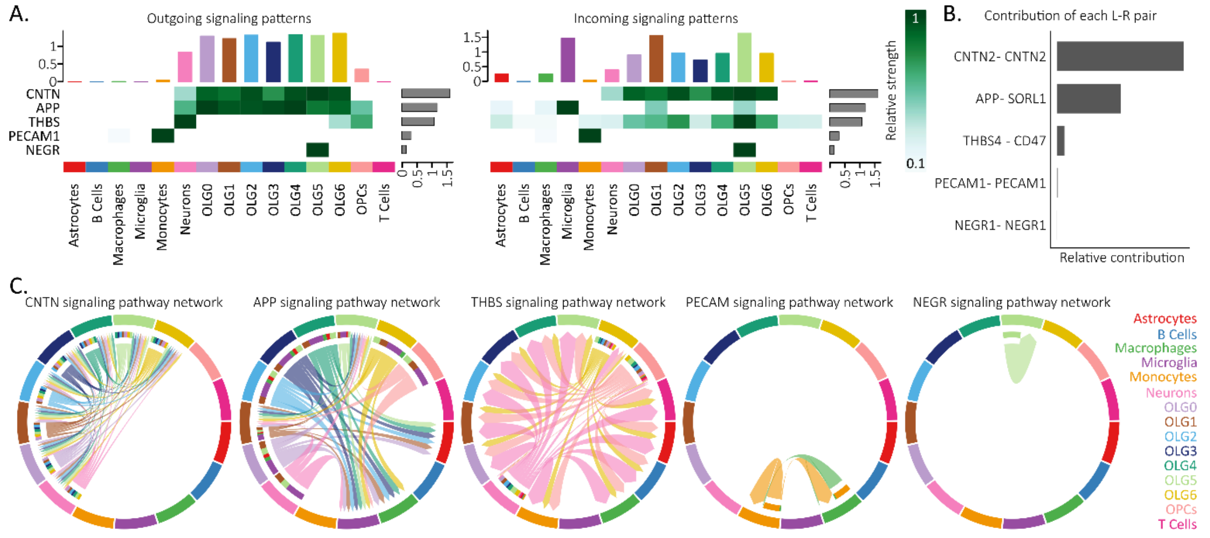


**Figure 5 Supplemental Legend. Intercellular communication networks in PD**

(A) Heatmap showing the summary of the signalling pathways that contribute to outgoing and incoming communication in PD. The colour bar represents the relative signalling strength of a signalling pathway across cell-types. The bars indicate the sum of the signalling strength of each cell-type per pathway. (B) Bar plot of relative contribution of each ligand-receptor pair to the signalling pathway. (C) Chord diagram showing the communication pattern of each signalling pathway in PD. L ligand; R receptor; OLG0-6 oligodendrocyte subtype 0-6.

**Supplemental Table**

**Table S1: Clinical metadata from patients**

PD = Parkinson’s disease; non-PD = clinical diagnosis other than Parkinson’s disease; HY = Hoehn and Yahr scale; UPDRS = Unified Parkinson's Disease Rating Scale; LED = levodopa equivalent dose;

| Project/ID | Diagnosis | Disease duration | HY | UPDRS off | UPDRS on | UPDRS difference | LED | Sex |
|---|---|---|---|---|---|---|---|---|
| PD_001 | PD | 17 | 4 | 23 | 18 | 5 | 1998 | M |
| PD_002 | PD | 3 | Na | 42 | 30 | 12 | 993 | M |
| PD_005 | PD | 11 | 2 | 31 | 8 | 23 | 855 | F |
| PD_006 | non-PD | 18 | Na | Na | Na | Na | Na | F |
| PD_007 | PD | 4 | 3 | 43 | 14 | 29 | 1075 | F |
| PD_008 | PD | 9 | 3 | 28 | 11 | 17 | 925 | M |
| PD_009 | PD | 17 | 2 | 23 | 8 | 15 | 990 | F |
| PD_011 | non-PD | 13 | Na | Na | Na | Na | Na | F |
| PD_012 | PD | 7 | 4 | 30 | 20 | 10 | 720 | F |
| PD_013 | non-PD | 9 | Na | Na | Na | Na | Na | F |
| PD_014 | non-PD | 21 | Na | Na | Na | Na | Na | M |
| PD_016 | PD | 8 | 3 | 27 | 8 | 19 | 600 | M |
| PD_017 | PD | 15 | 3 | 44 | 21 | 23 | 1098 | F |
| PD_018 | non-PD | 6 | Na | Na | Na | Na | Na | F |
| PD_020 | PD | 6 | Na | 15 | 9 | 6 | 798 | M |
| PD_021 | PD | 10 | Na | 69 | 45 | 24 | 800 | M |
